# Ion channel model reduction using manifold boundaries

**DOI:** 10.1101/2022.03.11.483794

**Authors:** Dominic G. Whittaker, Jiahui Wang, Joseph G. Shuttleworth, Ravichandra Venkateshappa, Jacob M. Kemp, Thomas W. Claydon, Gary R. Mirams

**Affiliations:** Centre for Mathematical Medicine & Biology, School of Mathematical Sciences, University of Nottingham, UK; Department of Biomedical Physiology and Kinesiology, Simon Fraser University, Burnaby, Canada

## Abstract

Mathematical models of voltage-gated ion channels are used in basic research, industrial and clinical settings. These models range in complexity, but typically contain numerous variables representing the proportion of channels in a given state, and parameters describing the voltage-dependent rates of transition between states. An open problem is selecting the appropriate degree of complexity and structure for an ion channel model given data availability. Here, we simplify a model of the cardiac human Ether-à-go-go Related Gene (hERG) potassium ion channel, which carries cardiac *I*_Kr_, using the manifold boundary approximation method (MBAM). The MBAM approximates high-dimensional model-output manifolds by reduced models describing their boundaries, resulting in models with fewer parameters (and often variables). We produced a series of models of reducing complexity starting from an established 5-state hERG model with 15 parameters. Models with up to 3 fewer states and 8 fewer parameters were shown to retain much of the predictive capability of the full model and were validated using experimental hERG1a data collected in HEK293 cells at 37°C. The method provides a way to simplify complex models of ion channels that improves parameter identifiability and will aid in future model development.

## 1 Introduction

Mathematical models of ion channel currents have been used for a wide variety of applications in cardiac research and drug discovery, with an increasing focus on making quantitative predictions for safety-critical applications [29]. However, these models usually contain numerous parameters and variables, which makes understanding their behaviour from the basic components challenging. The manifold boundary approximation method (MBAM) is a recently-developed method which constructs submanifold approximations of high-dimensional model manifolds at their boundaries [36], producing models with fewer parameters (and variables) whilst retaining much of the predictive capability of the original model. This reduction in complexity can improve parameter identifiability and offer greater insight into the connection between a model’s components and its output. The development of reduced models that are more practical to fit to experimental data may prove to be an important step towards cell- and patient-specific modelling.

The MBAM has been applied to a wide variety of model classes [36, 37], as well as action potential models within a cardiac modelling context [26, 17]. However, we believe it has yet to be applied directly to cardiac ion channel models, which may be another route to action potential model reduction. One disadvantage of applying the MBAM directly to action potential models is that it can lead to the complete removal of whole currents (such as *I*_Kr_ and *I*_Ks_ in [26]), which turns the model from a biophysically-detailed one to a semi-phenomenological one. Applying the MBAM to ion current models within action potential models offers the chance to reduce model complexity (and increase parameter identifiability) without necessarily sacrificing biophysical detail of a whole cell model.

Modelling the constituent ion channels/currents of the whole-cell cardiac electrical response is an active area of research with a rich history [31, 41]. However, a previous study revealed that the parameters in many models of cardiac ion channels are likely to be *unidentifiable* [16], which means it is not possible to determine their values uniquely by measuring model outputs. This is in conflict with the idea that a model’s structure and parameters provide insights into the underlying biophysical processes. The aim of this study was to create reduced models of the cardiac human Ether-àgo-go Related Gene (hERG) ion channel, a critical determinant of action potential repolarisation and focus of safety pharmacology [38], which retained the behaviour of the full model whilst containing fewer parameters and dynamic variables.

## 2 Materials and Methods

### 2.1 Manifold boundary approximation method

The manifold boundary approximation method (MBAM), first described in Transtrum and Qiu [36], is a model reduction algorithm which exploits the fact that many model outputs are bounded with a hierarchy of widths, a property which enables lower dimensional model approximations to be made at these boundaries. The *model manifold*, ℳ, can be thought of as an *N*-dimensional parameter space (*θ*_1_, *θ*_2_, …, *θ*_*N*_) manifold of a model embedded within an *M*-dimensional data space (the space of model output observations). In the data space the coordinates of the model manifold correspond to the system measurements (*y*_1_, *y*_2_, …, *y*_*M*_), with the point along the manifold closest to some desired data point representing the best model fit. In our case we chose a cost function which represented the model fit to reference system experimental data (see Section 2.4 for details), giving the cost function measurements of the full model, 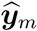, the parameters of which were fit to our own experimental data (see Section 2.4 for details), giving the cost function

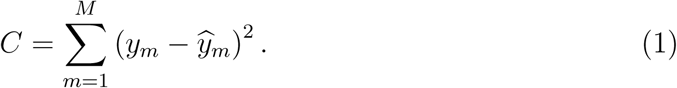

Many models in systems biology possess the property that certain parameters can take a wide range of values without greatly affecting the model output, which has been termed *sloppiness* [6, 40, 19]. The key to the MBAM lies in the fact that the model output space manifold can be extremely narrow in directions that are very sloppy, or in other words the model output does not change much even as you vary parameters (or combinations of parameters) to their plausible limits. This feature allows one to approximate the model with a manifold of reduced dimensionality by removing or combining parameters along a manifold boundary. We seek to reduce the dimensionality without greatly increasing the cost function value, and so from our starting point on the parameter manifold travel in the ‘sloppiest’ direction and then approximate the model along the boundary first encountered.

#### The MBAM proceeds as an iterative four-step algorithm with the following steps

1. The sloppy directions along ℳ are found by calculating the eigenvalues of an *N* × *N* matrix which has entries defined as

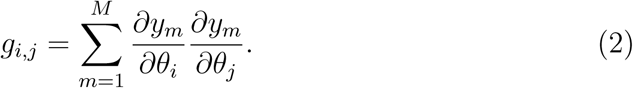

The eigenvector of **g** with the smallest eigenvalue, υ_0_, corresponds to the ‘sloppiest’ direction in parameter space.

2. In order to approach the manifold boundary, we use υ_0_ as an initial direction on ℳ and solve numerically a geodesic equation to find a path ***θ***(*τ*) through parameter space:

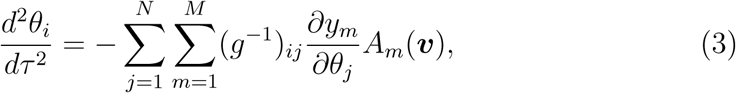

where υ = *d****θ****/dτ* and *A*_*m*_(υ) is the directional second derivative of *y*_*m*_ in the direction of the eigenvector υ:

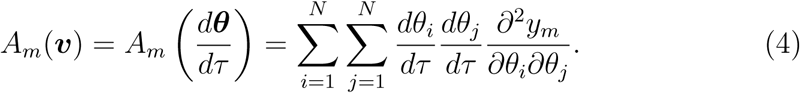

When the calculated path ***θ***(*τ*) approaches a boundary of ℳ, the smallest eigenvalue of **g** becomes small (much smaller than the next smallest) and approaches 0. This corresponds to a physically meaningful limit of the parameters in which either a single parameter or combinations of parameters are removed or combined to form new parameters.

For reasons of computational efficiency, we did not compute second order sensitivities directly, rather estimating *A*_*m*_(υ) using finite differences:

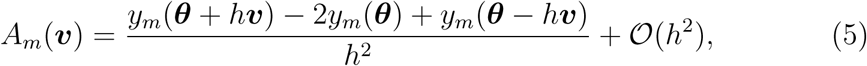

where *h* is the step size.

3. Deduce the reduced form of the model (which now has one fewer parameter). This model reduction step may be trivial, or may involve reformulation of model equations and removal of state variables.

4. Calibrate the new model with reduced parameter vector, ***θ***, to the full (original) model output by minimising the cost function, *C* (Equation 1).

An example of steps one and two of this procedure is shown in Figure 1 for the model we are going to introduce shortly (this example is the 4th iteration in Section 3.1 below, going from the r3 to r4 model). Although initially the ‘sloppiest’ eigenvector had components in multiple parameter directions, after following the geodesic path to a boundary the final eigenvector pointed exclusively in the direction of parameter 2 (Figure 1A), with the smallest eigenvalue approaching zero (Figure 1B). In a twodimensional slice of the parameter space, the geodesic can be seen to follow a canyon of the cost contour (Figure 1C), indicating that this path through parameter space incurs little to no change in model output. We note here that for steps 1 and 2 of the MBAM, all parameters were log-transformed to guarantee positivity, as suggested in [36, 37].

**Figure 1:**
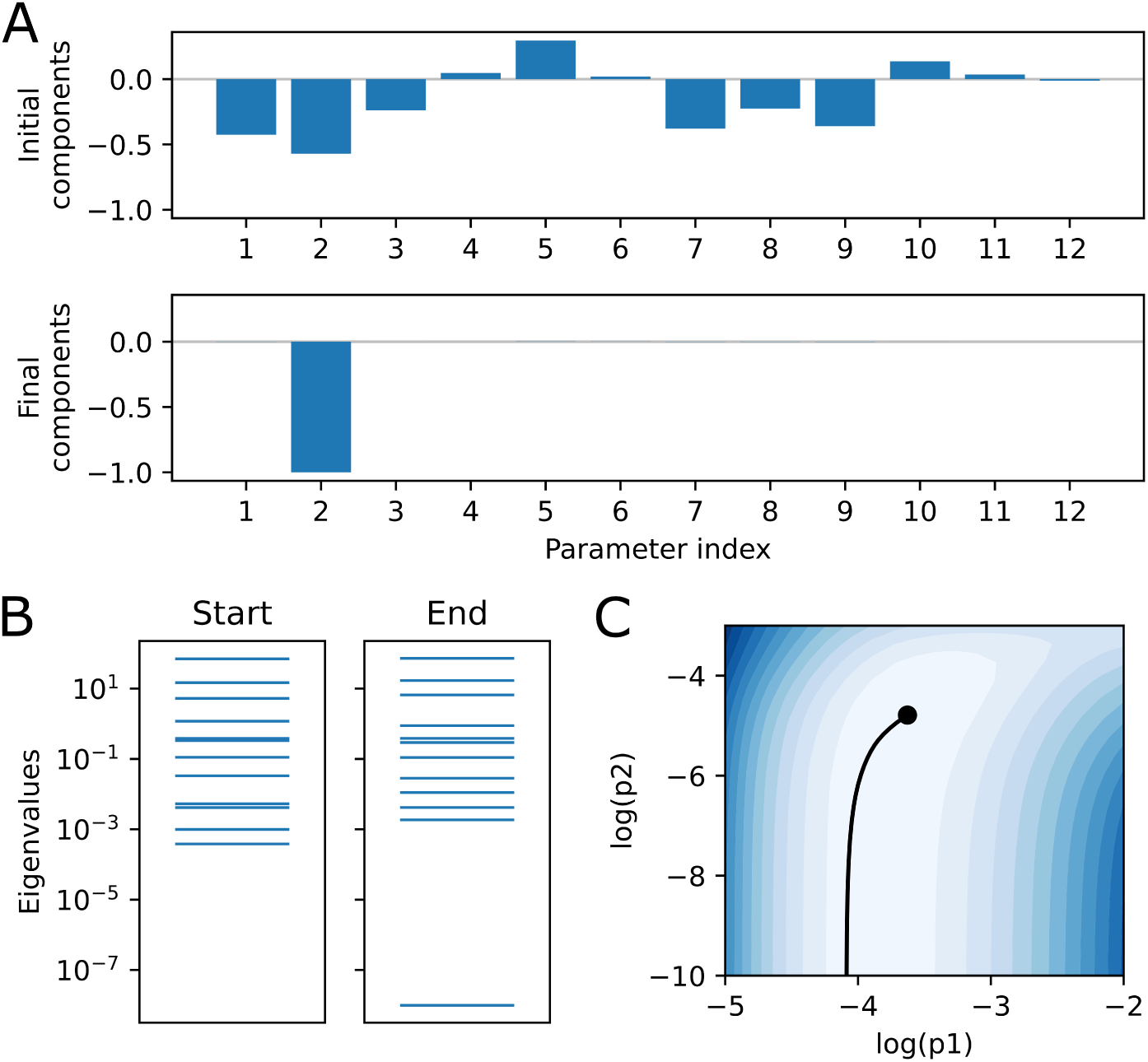
(A) Eigenvector components of the initial (sloppiest) and final parameter direction at the end of the geodesic path for an MBAM iteration (the 4th MBAM iteration in the results section for revision r3→ r4). (B) Eigenvalue spectra of **g** at the start and end of the geodesic path. (C) A plot of the geodesic path (black line) in a slice of log parameter space from the starting point denoted by a black circle. The plot is coloured according to evaluations of the cost function given in Equation 1, such that darker shades of blue represent worse agreement with the full model output.

We repeated this four-step process until the original model behaviour could not be reproduced within a reasonable error, which we defined using a mixed root mean square error [12], *e*_MRMS_, where

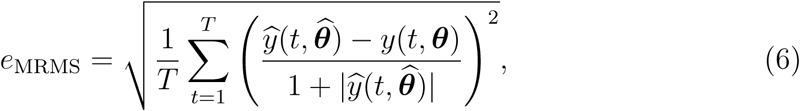

for an initial full model reference parameter vector 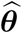 and *T* distinct time points (spaced 1 ms apart along the whole current trace). We chose an eMRMS threshold of 0.1 as the error beyond which a model could no longer satisfactorily reproduce the full model output.

Software to perform the MBAM was adapted from available python code provided by Dr Transtrum and colleagues (https://github.com/mktranstrum/MBAM); for a visual explanation of how the MBAM works, the interested reader is referred to a Michaelis Menten reaction kinetics toy model found in this repository and presented in detail in the Supplementary Information (see Supplementary Results and Supplementary Figure S1). In this work, python scripts were updated so that model equations are written as symbolic expressions using SymPy/SymEngine. All simulation codes and data pertaining to the MBAM and parameter inference described in Section 2.4 are freely available at https://github.com/CardiacModelling/model-reduction-manifold-boundaries.

### 2.2 Cardiac ion channel model

We used the well-established Wang model of hERG ion channel kinetics as our starting model [39]. This model contains 5 state variables (3 closed states, 1 open state, and 1 inactivated state) and 15 parameters (14 kinetic parameters governing the state transition rates and their voltage dependencies, and 1 conductance parameter). A schematic of the model is shown in Figure 2A, marked as revision zero, ‘r0’. If ***X*** = (*C*_1_, *C*_2_, *C* _3_, *O, I*) ^T^ is a vector of the state occupancies, the model is described by the system

**Figure 2:**
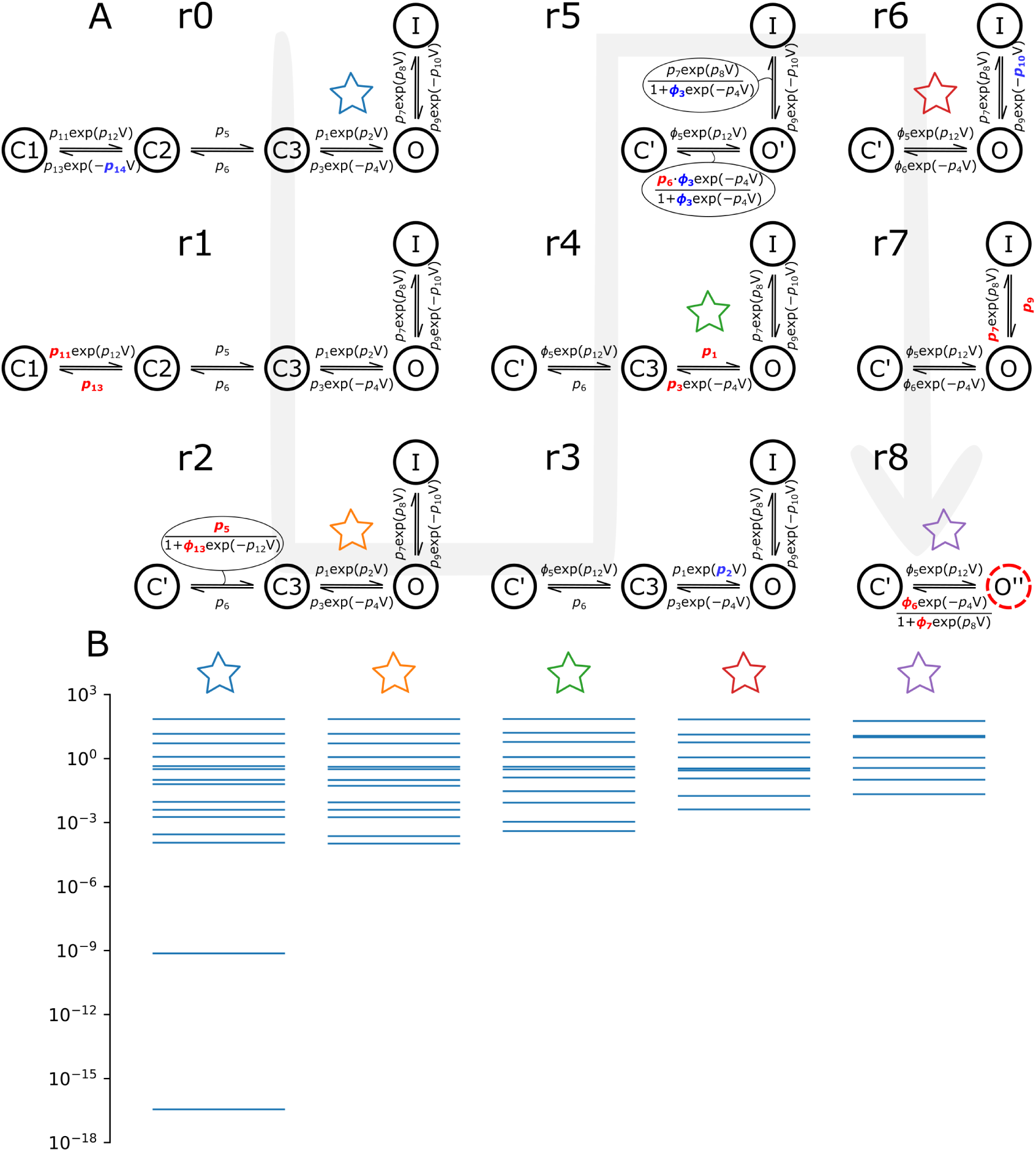
(A) Evolution of the Markov chain hERG model by Wang et al. [39] through subsequent iterations of the MBAM. Each structure shows a dynamic model (starting with the full model, r0) and the parameter changes which took place to get to the next reduced model, as in Table 1, guided by a grey arrow showing the direction of model reduction (from r0 to r8). As in Table 1, parameters highlighted blue → 0 and red → ∞ in the next reduction. The *O*′ and *O*″ states for the Wang-r5 and r8 models relate to the actual open probability through the relations *O* = *O*′ /(1+*ϕ*_3_ exp(−*p*_4_*V*)) and *O* = *O*″ /(1 + *ϕ*_7_ exp(*p*_8_*V*)), respectively. The dotted red line around the *O*″ state in the final reduction denotes that the maximal channel conductance. → ∞ (B) The eigenvalue spectra of **g** for selected models (denoted by coloured stars) under the shortened ‘staircase’ protocol described in Section 2.2.

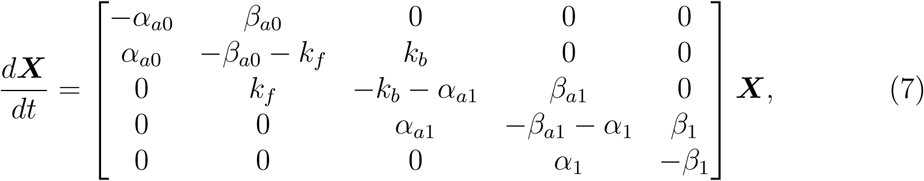

where

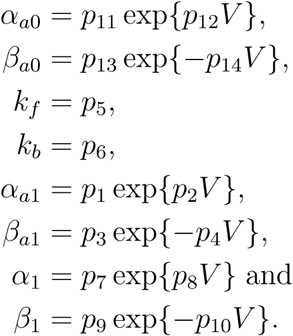

The current through the hERG channels is then given by

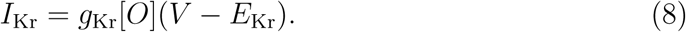

Here, *p*_1_, …, *p*_14_ represent kinetic parameters and *g*_Kr_ is the maximal conductance parameter; all are positive. In practice, we solve just four of the five ODEs, using the fact the probabilities sum to one to give *C*_1_ = 1 − (*C*_2_ + *C*_3_ + *O* + *I*). The initial parameters, 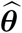, were obtained by fitting the 15 parameters of the model to data from our ‘staircase’ calibration protocol [23] at 37°C (see Section 2.4 for more details). The model output of interest (*y*_*m*_) used in Equations 1 and 2 was the current, *I*_Kr_, at time point *m*.

The input voltage protocol used for the MBAM procedure was a 5000 ms protocol which explored in a rapid way a similar range of voltages and time scales as our previously-published ‘staircase’ protocol [23], shown in Supplementary Figure S2. System measurements were made at 37 equally-spaced points which captured the main features of the current trace and appropriately weighted large, negative currents during which the channel is close to maximally open. It should be noted that in Equation 1 we used *M* = 37 for steps 1 and 2 of the MBAM algorithm and *M* = 5000 (1 ms spacing) when calibrating the model to the full model output in step 4.

Regarding the eigenvalue stopping criterion, a default value of 10^−6^ was used, which was typically sufficient to identify the geodesic limit with ease. This value occasionally required tuning as geodesic calculations can become very stiff when a boundary is approached (furthermore, ODE solver errors may result from parameters approaching infinity). Exact input settings including smallest eigenvalue thresholds used to generate the data in this study can be found in the Github repository (https://github.com/CardiacModelling/model-reduction-manifold-boundaries).

### 2.3 Electrophysiology experiments at 37^°^C

In order to test the predictive power of models reduced with the MBAM, we subsequently re-calibrated them to real experimental data. For parameter inference and model validation in this context we used HEK293 wild-type hERG1a expression system current traces at 37°C, which represented typical recordings from a previous study [22]. Briefly, HEK-293 cells cultured in DMEM supplemented with 10% FBS at 37°C with 5% CO_2_ were co-transfected with hERG1a in pcDNA3 and GFP in pcDNA3 using lipofectamine 3000 (Invitrogen). Cells were plated onto coverslips 14–16 h after transfection and cells with green fluorescence were selected for recordings. Whole cell patch clamp recordings were performed with an Axon Instruments 200B amplifier and Digidata 1440 A/D interface. Signals were acquired at a sampling frequency of 10 kHz and were filtered using a 4 kHz low pass Bessel filter.

During recordings, cells were superfused at 2 ml/min with solution containing (in mM): 140 NaCl, 4 KCl, 1.8 CaCl_2_, 1 MgCl_2_, 10 glucose, 10 HEPES (pH 7.4 with NaOH). Patch electrodes formed from borosilicate glass (Sutter Instruments) using a P-97 puller (Sutter Instruments) were filled with (in mM): 130 KCl, 1 MgCl_2_, 1 CaCl_2_, 10 EGTA, 10 HEPES, 5 Mg^2+^ATP (pH 7.2 with KOH). Electrodes had a resistance of 3.7–4.5 MΩ and series resistance was compensated 60–70%, without online leak subtraction, using the amplifier circuitry. The recording bath temperature was main-tained at 37°C using a TC-344B Warner Instruments temperature controller unit with bath chamber thermistor, heated platform, and inline perfusion heater. Upon whole cell formation, hERG1a current was recorded during a 2 s step to +20 mV followed by a step to − 65 mV (holding potential −80 mV) applied repeatedly at 0.2 Hz. Once peak tail current amplitude during the step to −65 mV stabilised, experimental recordings were undertaken.

### 2.4 Parameter inference using real data

As described previously [22], maximum likelihood estimation was used to infer model parameters from the experimental data, by constructing a likelihood function based on independent and identically distributed Gaussian noise on each data point:

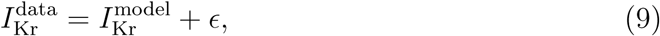

where ϵ 𝒩 (0, *σ*^2^) [22]. Under this scheme, the log-likelihood of a given set of parameters is proportional to

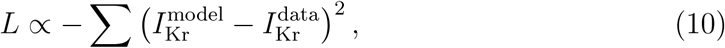

where we sum over the time points in the current trace for the calibration protocol data. The most likely parameter set is thus identical to that given by a least-sum-ofsquare-errors fit as used in our cost function *C* here, Equation 1.

All fitting used a Myokit [8] model in PINTS [10], using the CVODE solver [11] with absolute and relative tolerances of 10^−8^ and maximum time step of 0.1 ms. We used CMA-ES with 50 repeats from different initial guesses for optimisation to real data, and as in Kemp et al. [22] the optimiser worked with log-transformed parameters for those that are non–voltage-dependent in the transition rates [9, 41].

## 3 Results

### 3.1 A series of reduced models

A summary of model reductions at each iteration of the MBAM and the *e*_MRMS_ of the associated reduced model is given in Table 1. Figure 1 showed an example of the progress of the algorithm from r3 to r4 as it establishes which parameters will be reduced as the boundary is approached. The first 9 iterations only are shown in Table 1, as after this the reduced model exceeded our *e*_MRMS_ threshold. We can see that 4/9 of the reductions involved a single parameter tending to 0, 4/9 of the reductions involved two parameters tending to infinity, the finite ratio of which formed a new parameter, and 1 reduction involved one parameter tending to zero and another tending to infinity, the finite product of which formed a new parameter. A detailed breakdown of each MBAM iteration and the effect it had on model equations is given in the Supplementary Information.

Figure 2A shows model structures for the full model and the series of models of reduced complexity obtained with the MBAM. It should be noted that the Wang-r5 and Wang-r8 models have extra voltage-dependence in the expressions for the open probability which cannot be represented in the Markov chain diagrams. In the case of the Wang-r8 model, this corresponds to instantaneous inactivation, thus giving the model more flexibility than suggested by the simple closed-open model structure. The spread of eigenvalues of **g** (Equation 2) for selected models is presented in Figure 2B. From this we can see that the eigenvalue spectrum of reduced models spanned fewer orders of magnitude than the full model, with the spread decreasing with the level of model reduction, signifying reduced *sloppiness* of the parameter sensitivities.

**Table 1:**
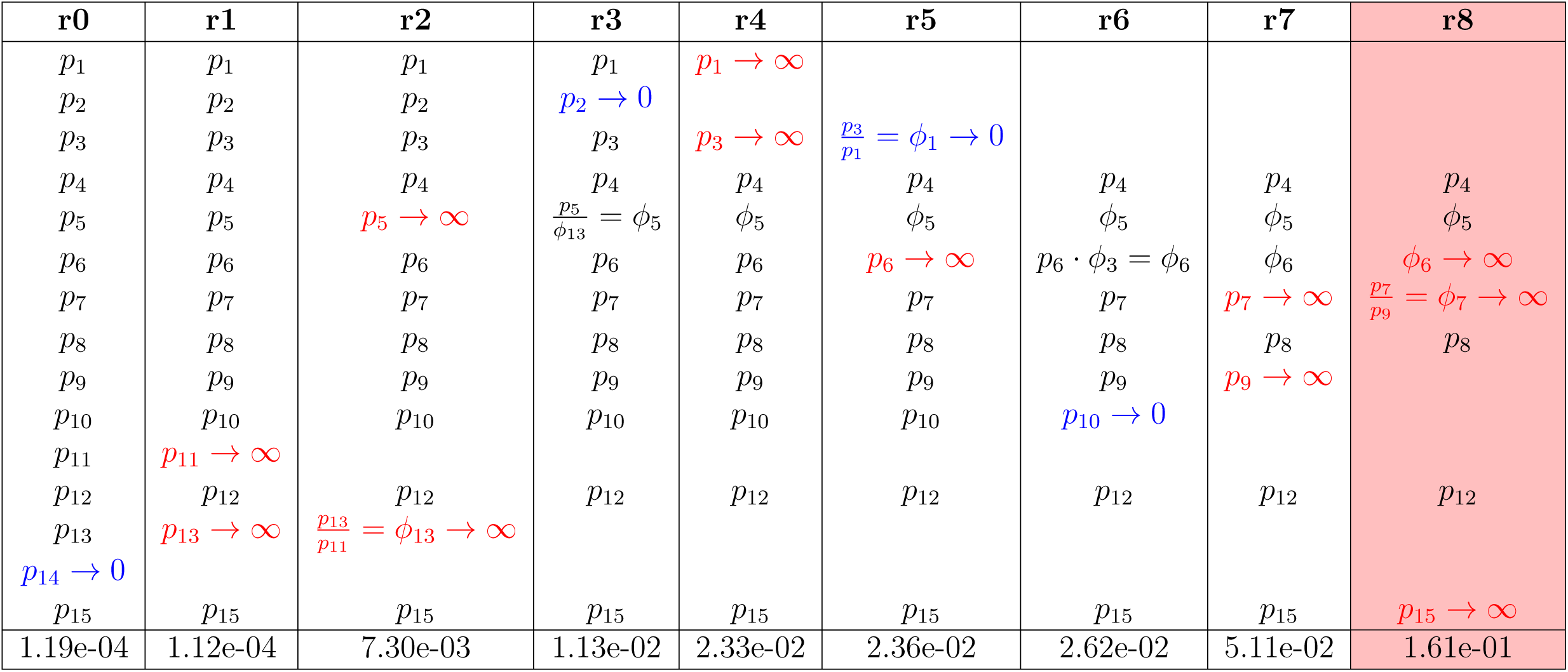
A table showing parameter changes between iterations of the MBAM reduction algorithm. Each column shows a model (starting with the full model, r0) and the changes which took place to get to the next reduced model. Parameters highlighted in blue → 0 and red → ∞. In some cases, parameters were combined to form new parameters. The bottom row shows the calculated *e*_MRMS_. The rightmost column is highlighted in red as it exceeded our threshold eMRMS of 0.1.

### 3.2 Model reduction with the MBAM improves parameter identifiability

In order to demonstrate the advantage of performing model reduction, we next performed an exercise in which the parameters of all reduced models were fit to our WT hERG channel current trace at 37°C (as was done to obtain the initial parameter set, 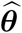, in the full model). This process was repeated 50 times for each model, sampling from the parameter search space each time to give a different initial guess. The best 30 parameter sets for the full model are plotted in Figure 3A. The results reveal that many of the parameters could take on a wide range of values which spanned several orders of magnitude whilst still giving a model output consistent with the experimental data. To illustrate this point further, Figure 3C shows two parameter sets with huge differences in the values of many parameters which produce highly similar model outputs in response to the same input (Figure 3D). This tells us that the parameters in this model are *practically unidentifiable* for this particular experiment. It is important to stress that practical identifiability is a property of both model and experiment — given that the parameters of our full model are not *a priori* unidentifiable, we could in theory design a new experiment which would enable us to determine uniquely the values of all parameters (given structural identifiability) [16, 41]. Indeed, the original Wang et al. [39] model developers did use different experimental data and carefully considered a range of structures when motivating this choice of model [39].

**Figure 3:**
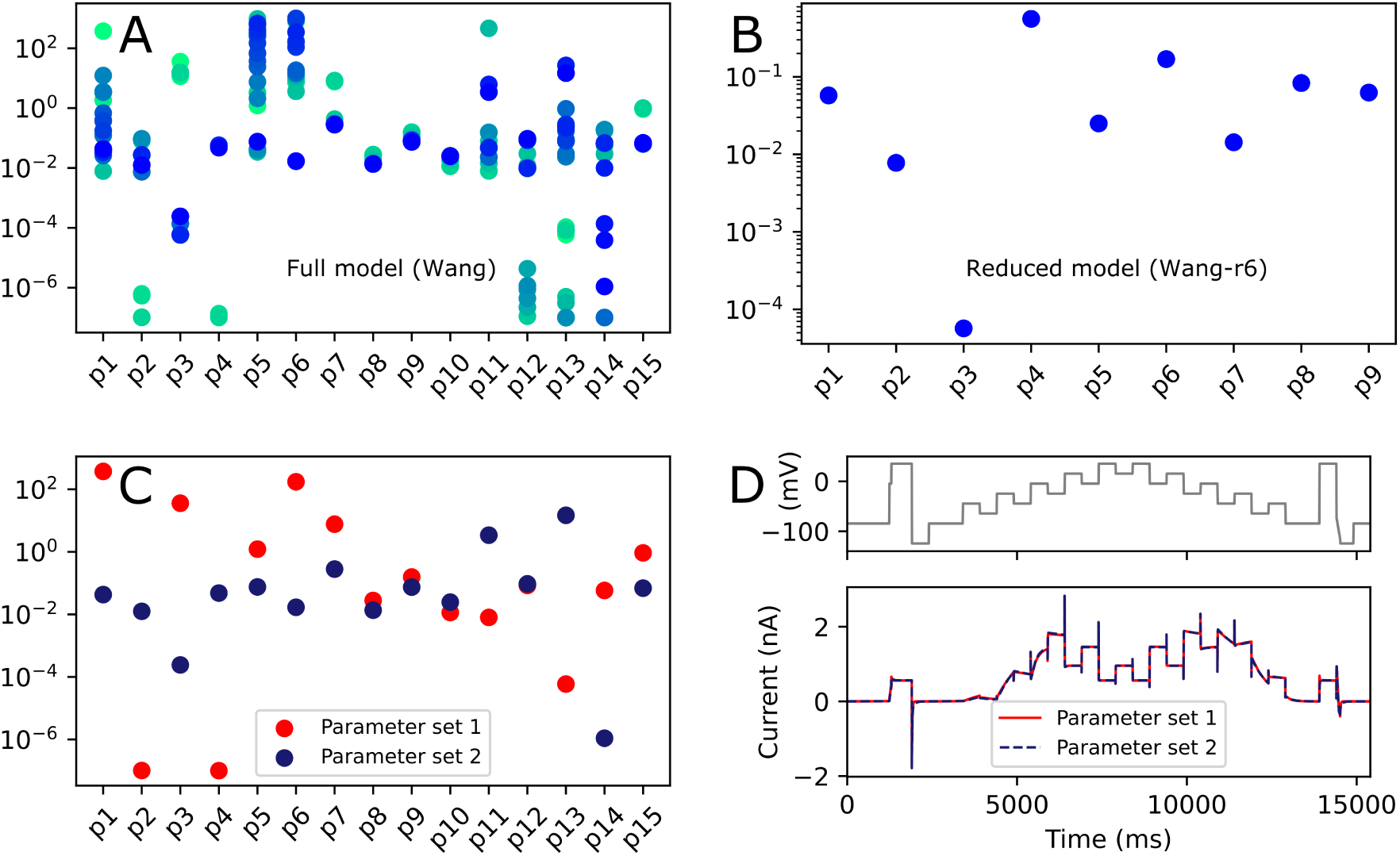
A practical assessment of model identifiability. Plotted are the inferred parameter values in (A) the full Wang model [39] and (B) a reduced model (Wang-r6) for 50 repeats of fitting from different initial guesses to experimental ‘staircase’ calibration protocol hERG channel currents at 37°C [22]. The 30 best parameter sets are shown in each case, from lowest (green) to highest (blue) likelihood. Note that in panel (B) the 30 inferred parameter sets are overlapping. (C) Two parameter sets of the full Wang model which show large divergence for many parameters but are both consistent with the experimental data, giving highly similar model outputs in response to the same input voltage protocol, as shown in (D).

Figure 3B shows the results of our repeated parameter fitting exercise for a model which was reduced through 6 iterations of the MBAM (termed the Wang-r6 model). We focus on this model as it is the first model with fully identifiable parameters from the experiment (Supplementary Figure S3) whilst also being the most reduced model with simple, biophysically-interpretable rates of the form *A* · exp(*B* · *V*) on each of the transitions. We can see in this case that we have convergence of our parameter estimates — all of our inferred values in the best 30 parameter sets occupy a very small (overlapping) region of parameter space. This tells us that the parameters in our reduced model are *practically identifiable* for this particular experiment, putting us in a stronger position to to draw conclusions about the underlying biophysical processes.

### 3.3 Reduced models retain high predictive capability

As described in the previous section, after reducing the Wang model through several iterations using the MBAM we fit the parameters of the new, reduced models to ‘staircase’ calibration protocol experimental data (convergence of parameter estimates from different initial guesses is shown for the Wang-r6 model in Figure 3B and for all models in Supplementary Figure S3). Focusing again on the Wang-r6 model and also the Wang-r8 model (the most reduced model with acceptable error), the close correspondence between model and experiment achieved in model calibration is shown in Figure 4A. Furthermore, the calibrated reduced models excellently predicted the response to a wealth of ‘unseen’ validation data collected in the same cell [22]. Specifically, the Wang-r6 and Wang-r8 reduced models predicted with quantitative accuracy the response to a complex series of cardiac action potential waveforms (Figure 4B) [5] and shortened versions of traditional activation and inactivation voltage protocols (Figure 4C) plus associated summary data (Figure 4D) with highly similar model output to the full Wang model. The only very noticeable area of model discrepancy was in the time constant of deactivation at higher voltages. This is due to the fact that the structures of the Wang-r6 and Wang-r8 reduced models can only produce one time course of deactivation (where at least two exist in the experimental data). However, we can see from Figure 4B that this is not an important feature of the model for making predictions of resurgent hERG currents in a physiologically-relevant context-of-use.

**Figure 4:**
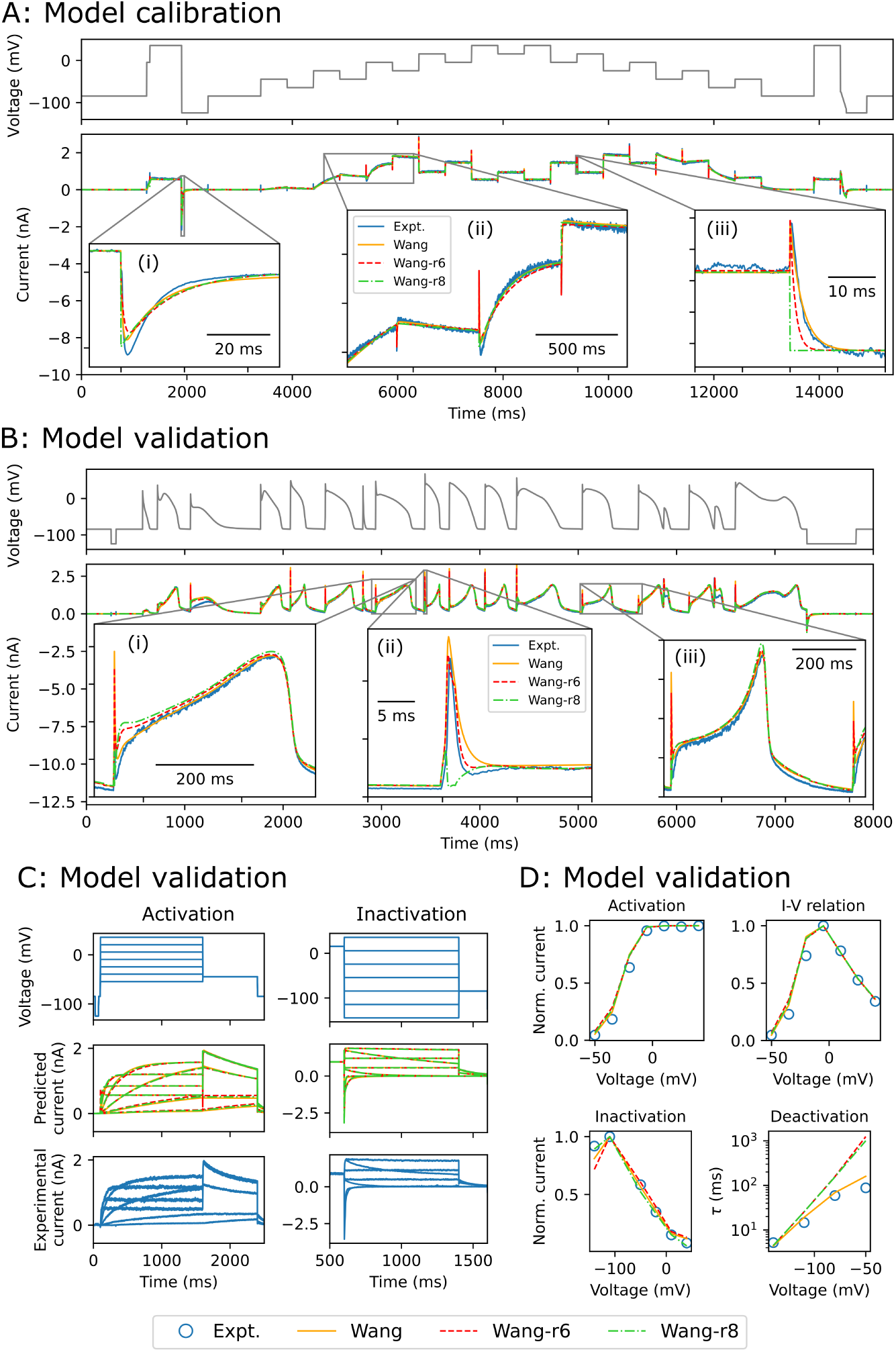
(A) A comparison of the full Wang model and Wang-r6/Wang-r8 reduced model fits to experimental data under the ‘staircase’ calibration protocol. (B) Prediction of the full Wang model and Wang-r6/Wang-r8 reduced models under a complex action potential waveform validation protocol. (C) Predictions of the full Wang model and Wang-r6/Wang-r8 reduced models under shortened versions of traditional activation and inactivation protocols. (D) Comparisons of summary data between the full Wang model and Wang-r6/Wang-r8 reduced models and experiment corresponding to the data shown in (C). All experimental data were recorded in HEK293 cells at 37°C [22] (see Section 2.3 for details).

Fits to the ‘staircase’ protocol for all models are shown in Supplementary Figure S4, from which it can be seen that reducing the model through as many as 8 iterations of the MBAM had only a very small effect on the ability of the models to fit the data. Similarly, all reduced models were able to predict the response to a complex series of cardiac action potential waveforms (Supplementary Figure S5) and shortened versions of traditional activation and inactivation protocols (Supplementary Figure S6) with similar accuracy as the full model. The only noticeable exceptions were that the Wang-r7 and Wang-r8 models underestimated the amplitude of hERG transient currents under the complex action potential validation protocol (Supplementary Figure S5), and all models reduced beyond the Wang-r3 model exhibited appreciable discrepancy in the time constant of deactivation (Supplementary Figure S6), which we revisit in the Discussion.

## 4 Discussion

### 4.1 Main findings

Our study demonstrates that the MBAM is a viable approach to reducing cardiac ion channel models. We reduced the established Wang et al. [39] model of the critically important cardiac hERG channel from one with 15 parameters and 5 variables to a series of reduced models with as few as 7 parameters and 2 variables which preserved the predictive capability of the full model. This was demonstrated to improve practical identifiability of the parameters, as shown through convergence of parameter estimates when fitting to experimental data from different initial guesses (Figure 3B and Supplementary Figure S3). Another way of framing this is that we reduced the ‘sloppiness’ of the parameter sensitivities, as shown by the smaller spread of eigenvalues in Figure 2B. It has been suggested that *sloppiness*, in which well-constrained predictions can arise from poorly-constrained parameters, is a “universal” property of systems biology models [6, 40, 19]. However, others have pointed out that this sloppiness is simply unidentifiability which can be rectified through novel experimental design, i.e. by performing the ‘right’ experiments [3], whereas others have proposed that sloppiness and lack of identifiability are not equivalent [7]. Using the definition of a sloppy model provided by Chis et al. [7], i.e. that λ_min_/λ_max_ 10^−3^, our three most reduced models (Wang-r6, Wang-r7 and Wang-r8) would be considered sloppy yet identifiable. We demonstrated convergence of parameters from different initial guesses for each of these models (Supplementary Figure S3), suggesting that the model complexity and informativeness of the experiment are appropriately matched — we would, therefore, favour identifiability criteria over those pertaining to sloppiness, in line with the conclusions of that study [7].

Using previously-reported experimental data at 37°C [22], we showed that models reduced with the MBAM retained a large amount of flexibility and predictive power of the full model (Supplementary Figures S5 and S6). Not only were the reduced models able to fit the calibration data very well (this was partially by design, as the calibration voltage protocol was highly similar to the voltage input protocol used to generate the system measurements for the MBAM), they were also able to predict a vast amount of independent, ‘unseen’ validation data recorded from the same cell (e.g. see Wang-r6 and Wang-r8 model predictions in Figure 4). Especially impressive is the fact that the reduced models were found to be highly predictive in the context of a complex, physiologically-relevant series of cardiac action potential waveforms which explores channel dynamics under both normal and abnormal action potential morphologies, including delayed- and early-afterdepolarisations [5].

The most noticeable area of model discrepancy (or model *mismatch* — the interested reader is referred to Table 1 in Lei et al. [24] for a list of equivalent terminologies for inverse problem concepts) was in the time constant of deactivation (Figure 4D) extracted from the inactivation protocol validation data (Figure 4C). Supplementary Figure S6 highlights that the reduction which took place between the Wang-r3 and Wang-r4 model in particular increased the divergence between model and experiment. This step, illustrated in Figure 1, corresponded to the parameter *p*_2_→0, reducing the model from one with two voltage-dependent deactivation transition rates to a model with two deactivation transition rates, only one of which is voltage dependent. Following this reduction the model was no longer able to deactivate following the biexponential time course seen in the data, hence the greater discrepancy. Nonetheless, models reduced past this point preserved the steady state channel kinetics well, and this feature of the model was shown to be unimportant for making predictions of resurgent hERG currents under physiologically-relevant AP waveforms (Figure 4B), although the two most reduced models underestimated the amplitude of hERG transient currents (Supplementary Figure S5).

Model selection for ion channel models remains a challenging and unresolved problem [28, 35, 27]. Starting from an existing, established model in the literature, we showed that ion channel model reduction by the MBAM can offer insights into the underlying biophysical processes by reducing and refining the structure and parameters of a model, thus aiding in the model selection process. Rather than trying to select from a large range of available models in the literature, we demonstrated that the MBAM can be used to distil the components of an existing model which are necessary to give a predictive model. In our case, we demonstrated that removing two of the three closed states present in the original model of hERG channel kinetics described by Wang et al. [39] resulted in models which retained predictive accuracy. Whilst we are not claiming that the structure of any of our reduced models (such as the Wang-r6 model) give a more accurate representation of the true underlying molecular reality of the channel, we do suggest that these fundamental components of the system can explain a lot of the channel dynamics at 37°C and are thus sufficient to form the basis of a predictive and well-parameterised mathematical model. Interestingly, Di Veroli et al. [13] also settled on a simpler, single time constant of activation/deactivation representation of hERG channel dynamics at 37°C compared to their model at room temperature, which could produce two time courses of activation/deactivation.

### 4.2 Relation to previous work and future outlook

The relationship between the MBAM and other reduction techniques has been discussed in detail previously [36, 37]. As outlined in this paper, cardiac ion channel models typically contain large numbers of parameters relating to transition rates between numerous closed, open, and inactivated states, which may result in parameter unidentifiability [16] and divergence in predictions between models of the same channel kinetics [5, 1]. Multiple plausible models also means it is difficult to understand the relationship between model outputs and model parameters, suggesting there is a need for ion channel models of reduced complexity. Whilst this problem is wellknown when it comes to models of the cardiac action potential [42, 20], with several reduced/minimal models having been created already [2, 15, 30] (including through use of the MBAM [26, 17]), considerably less has been done in terms of reducing their constituent ion channel sub-models (more of a concern for Markov chain models than simpler Hodgkin-Huxley formulations).

Reducing the ion channel sub-models within full cellular action potential models has the advantage of preserving biophysical detail and therefore drug or mutation targets whose effects we may wish to model. For example, within a 67 variable myocyte model Ariful Islam et al. [4] reduced a 13-state sodium channel Markov model and 10-state potassium channel Markov model to two-gate Hodgkin-Huxley (HH) models. That work relied on using approximate bisimulation between the full Markov models and two-state HH invariant manifold reductions of the Markov channel dynamics [21]. Other model reduction techniques which have been applied to ion channel models include combining states (‘lumping’) and fast/slow analysis to separate time scales [18, 34, 33]. To decide which states to combine in lumping approaches, intuition of the modelling problem may be used, or each possible choice may be evaluated as in the ‘Proper Lumping’ technique [14]. In contrast, the MBAM seeks only to reduce the number of parameters whilst having little impact on outputs. The MBAM may lump states, as it did in r1→ r2 and r4→ r5 in our reductions. However, the MBAM is more flexible in the sense that the resulting reduction in number of parameters may be associated with model reductions that do not use lumping, as we saw above for most of our hERG model reductions, and has the benefit of semi-automatically suggesting the next reduction based on sensitivities rather than having to exhaustively try all combinations of states.

Models of the cardiac hERG ion channel are frequently used in the simulation of genetic mutations and drug effects, due to the medical and pharmaceutical relevance of hERG-related abnormalities. Some additional consideration is therefore warranted regarding how this method might be applied in these contexts. Regarding state-specific drug block, the method allows one to choose which observations to use to guide the model reduction. Accordingly, if we have a trusted complex model, it would be possible to preserve both the open and inactivated state occupancies, which would ensure the model remains relevant for use in conjunction with existing models of drug kinetics, which for hERG typically include binding to only open and/or inactivated states (e.g. [25]). As for genetic mutations, applying the MBAM separately for each mutant would produce models which are able to shed light on how a mutation affects the channel, with no requirements to be defined *a priori*.

An approach to parameter identification which circumvents the need for model reduction altogether is to fix the values of certain parameters based on experimental estimates or inheritance from previous models, fitting only the remaining parameters in the model [37]. This approach is relatively common not just in the field of cardiac modelling, but also in the relatively new discipline of quantitative systems pharmacology. Whilst this does reduce the dimensionality of the parameter search space, it does not make the model conceptually simpler or necessarily help to illuminate the connection between model parameters and output, unlike model reduction methods such as the MBAM. The MBAM may therefore also be of great utility in this context, in which the desirability of models with identifiable parameters has begun to be appreciated [32].

## 5 Conclusions

To conclude, we have demonstrated the viability of using the manifold boundary approximation method to reduce models of ion channels whilst retaining a high level of predictive power. This approach is a very promising way to simplify ion channel models whilst improving parameter identifiability. It maintains a strong connection between the biophysically-based model parameters, states and outputs from complex models and the same properties within algorithmically-derived simplified models.

## Supporting information

Supplementary Results

## Acknowledgements

This work was supported by the Wellcome Trust (grant no. 212203/Z/18/Z). GRM and DGW acknowledge support from the Wellcome Trust via a Wellcome Trust Senior Research Fellowship to GRM. This research was funded in whole, or in part, by the Wellcome Trust [212203/Z/18/Z]. For the purpose of open access, the author has applied a CC-BY public copyright licence to any Author Accepted Manuscript version arising from this submission.

## Notes

https://github.com/CardiacModelling/model-reduction-manifold-boundaries

### Competing Interest Statement

The authors have declared no competing interest.

### Summary of Updates

Figure 4A and B has been updated to show more detail on short 'inactivation spike' currents. The discussion has been expanded to contrast with the 'Proper Lumping' technique and discuss how the method might be used in modelling drug binding to ion channels.

