## Supplementary Results for "Ion channel model reduction using manifold boundaries"

### Ion channel model reduction using manifold boundaries – Supplementary Information

May 16, 2022

#### 1 Supplementary Results

##### 1.1 Toy model

In order to better explain how the manifold boundary approximation method (MBAM) works, we here show application of the method to a simple, "toy model" problem, which is available at `https://github.com/mktranstrum/MBAM` and [2]. We consider the dynamics of the Michaelis Menten reaction:

$$\dot{x}(t) = \frac{-\rho_1 x(t)}{\rho_2 + x(t)}, \tag{1}$$

where $\rho_1$ and $\rho_2$ are parameters of the model, and the system output is measured at three time points.

A single iteration of the MBAM shows that both parameters tend to infinity along the geodesic path (Supplementary Figure S1A). Plotting the geodesic path in parameter space illustrates how the geodesic path follows a canyon of the cost contour, i.e. a path towards the manifold boundary in which the value of the cost function does not increase greatly (Supplementary Figure S1B). Using the same colour scheme, we can plot the path of the geodesic along the manifold (in the "data" space of the three system observations) towards the boundary which corresponds to both parameters tending to $\infty$ (Supplementary Figure S1C). Evaluating the model at this limit, we see that the denominator $\approx \rho_2$ as $\rho_2 \gg x(t)$, and so we are left with a linear model

$$\dot{x(t)} = -\phi x(t), \tag{2}$$

where $\phi$ is a new parameter formed by the finite ratio $\rho_1/\rho_2$.

To summarise the MBAM and how it applies to this particular example:

- There are sloppy directions in parameter space in which parameter values change without changing the cost function value appreciably, as shown in Supplementary Figure S1A and B
- This trajectory in parameter space corresponds to moving along the shortest width of the model manifold (Supplementary Figure S1C)
- Eventually a limit is encountered in which a parameter/combination of parameters $\to 0/\infty$ and the model can be reformulated to one with fewer parameters (and sometimes variables), as in Equation 2
- The MBAM calculates a geodesic path along the model manifold in sloppy directions of parameter space as a way of reducing a model without changing significantly its model output and thus cost function value (Supplementary Figure S1D)

For a further step-by-step description and practical application of the MBAM with provided code (in Matlab) see `https://pengqiu.gatech.edu/software/model_manifold/html/publish_MBAM_example.html`.

#### 1.2 Iterations of the MBAM – hERG channel model

The system equations of the Wang model [3] are:

$$\dot{C2} = p_{11}\exp(p_{12}V)C1 + p_6C3 - (p_{13}\exp(-p_{14}V) + p_5)C2, \tag{3}$$

$$\dot{C3} = p_5C2 + p_3\exp(-p_4V)O - (p_6 + p_1\exp(p_2V))C3, \tag{4}$$

$$\begin{aligned}\dot{O} = {} & p_1\exp(p_2V)C3 + p_9\exp(-p_{10}V)I - \\ & (p_3\exp(-p_4V) + p_7\exp(p_8V))O,\end{aligned} \tag{5}$$

$$\dot{I} = p_7\exp(p_8V)O - p_9\exp(-p_{10}V)I, \tag{6}$$

where we have removed the ODE for $C1$ using the constraint that

$$C1 + C2 + C3 + O + I = 1. \tag{7}$$

##### 1.2.1 Iteration 1

The first iteration corresponded to the boundary where $p_{14} \to 0$. As this parameter appears in the rate $p_{13}\exp(-p_{14}V)$, we can simply write this rate as $p_{13}$, giving the new system equations:

$$\dot{C2} = p_{11}\exp(p_{12}V)C1 + p_6C3 - (p_{13} + p_5)C2, \tag{8}$$

$$\dot{C3} = p_5C2 + p_3\exp(-p_4V)O - (p_6 + p_1\exp(p_2V))C3, \tag{9}$$

$$\begin{aligned}\dot{O} = p_1\exp(p_2V)C3 + p_9\exp(-p_{10}V)I- \\ (p_3\exp(-p_4V) + p_7\exp(p_8V))O,\end{aligned} \tag{10}$$

$$\dot{I} = p_7\exp(p_8V)O - p_9\exp(-p_{10}V)I, \tag{11}$$

##### 1.2.2 Iteration 2

The second iteration corresponded to the boundary where $p_{11}, p_{13} \to \infty$. Dividing Equation 8 by $p_{11}$ gives us

$$C1 = \phi_{13}\exp(-p_{12}V)C2,$$

where $\phi_{13} = p_{13}/p_{11}$. As C1 can now be expressed as a function of C2, we lump them together into a new state, $C' = C1 + C2$, giving us

$$\dot{C'} = p_6C3 - p_5C2 = p_6C3 - \frac{p_5}{1 + \phi_{13}\exp(-p_{12}V)}C'.$$

This reduces the number of variables in the system by one, giving the new, reduced system equations:

$$\begin{aligned}\dot{C3} = \frac{p_5}{1 + \phi_{13}\exp(-p_{12}V)}C' + p_3\exp(-p_4V)O- \\ (p_6 + p_1\exp(p_2V))C3,\end{aligned} \tag{12}$$

$$\begin{aligned}\dot{O} = p_1\exp(p_2V)C3 + p_9\exp(-p_{10}V)I- \\ (p_3\exp(-p_4V) + p_7\exp(p_8V))O,\end{aligned} \tag{13}$$

$$\dot{I} = p_7\exp(p_8V)O - p_9\exp(-p_{10}V)I, \tag{14}$$

with

$$C' + C3 + O + I = 1.$$

##### 1.2.3 Iteration 3

The third iteration corresponded to the boundary where $p_5, \phi_{13} \to \infty$. As $\phi_{13} \gg 1$, the $C' \to C3$ transition $\approx \frac{p_5}{\phi_{13}\exp(-p_{12}V)}$, which we write as $\phi_5 \exp(p_{12}V)$. The new system equations are thus:

$$\begin{aligned}\dot{C3} = \phi_5 \exp(p_{12}V)C' + p_3 \exp(-p_4 V)O - \\ (p_6 + p_1 \exp(p_2 V))C3,\end{aligned} \tag{15}$$

$$\begin{aligned}\dot{O} = p_1 \exp(p_2 V)C3 + p_9 \exp(-p_{10}V)I - \\ (p_3 \exp(-p_4 V) + p_7 \exp(p_8 V))O,\end{aligned} \tag{16}$$

$$\dot{I} = p_7 \exp(p_8 V)O - p_9 \exp(-p_{10}V)I, \tag{17}$$

with

$$C' + C3 + O + I = 1.$$

##### 1.2.4 Iteration 4

The fourth iteration corresponded to the boundary where $p_2 \to 0$. As this parameter appears in the rate $p_1 \exp(p_2 V)$, we can simply write this rate as $p_1$, giving the new system equations:

$$\dot{C3} = \phi_5 \exp(p_{12}V)C' + p_3 \exp(-p_4 V)O - (p_6 + p_1)C3, \tag{18}$$

$$\begin{aligned}\dot{O} = p_1 C3 + p_9 \exp(-p_{10}V)I - \\ (p_3 \exp(-p_4 V) + p_7 \exp(p_8 V))O,\end{aligned} \tag{19}$$

$$\dot{I} = p_7 \exp(p_8 V)O - p_9 \exp(-p_{10}V)I, \tag{20}$$

with

$$C' + C3 + O + I = 1.$$

##### 1.2.5 Iteration 5

The fifth iteration corresponded to the boundary where $p_1, p_3 \to \infty$. Dividing Equation 18 by $p_3$ gives us

$$C3 = \phi_3 \exp(-p_4 V)O, \tag{21}$$

where $\phi_3 = p_3/p_1$. Due to the fact that O can be expressed as a function of C3, we consider a new state, $O'$, which is equal to $O + C3$. This gives us

$$\dot{O'} = \phi_5 \exp(p_{12}V)C' + p_9 \exp(-p_{10}V)I - p_6 C3 - p_7 \exp(p_8 V)O,$$

where

$$C3 = \frac{\phi_3 \exp(-p_4 V)}{1 + \phi_3 \exp(-p_4 V)} O',$$
$$O = \frac{O'}{1 + \phi_3 \exp(-p_4 V)}.$$

This again reduces the number of variables in the system by one, giving the new, reduced system equations:

$$\begin{aligned} \dot{O}' = \phi_5 \exp(p_{12} V) C' + p_9 \exp(-p_{10} V) I - \\ \left( p_6 \left( \frac{\phi_3 \exp(-p_4 V)}{1 + \phi_3 \exp(-p_4 V)} \right) - \frac{p_7 \exp(p_8 V)}{1 + \phi_3 \exp(-p_4 V)} \right) O' \end{aligned} \tag{22}$$

$$\dot{I} = \frac{p_7 \exp(p_8 V)}{1 + \phi_3 \exp(-p_4 V)} O' - p_9 \exp(-p_{10} V) I, \tag{23}$$

with

$$C' + O' + I = 1.$$

In this case we solve the ODEs as normal, but remember that the actual open probability is given by $O = O'/(1 + \phi_3 \exp(-p_4 V))$.

##### 1.2.6 Iteration 6

The sixth iteration corresponded to the boundary where $\phi_3 \to 0$ and $p_6 \to \infty$. From this we can see that the denominator $1 + \phi_3 \exp(-p_4 V) \to 1$ and we create a new parameter $\phi_6 = \phi_3 \cdot p_6$. The new system equations are thus given by

$$\begin{aligned} \dot{O} = \phi_5 \exp(p_{12} V) C' + p_9 \exp(-p_{10} V) I - \\ \left( \phi_6 \exp(-p_4 V) - p_7 \exp(p_8 V) \right) O, \end{aligned} \tag{24}$$

$$\dot{I} = p_7 \exp(p_8 V) O - p_9 \exp(-p_{10} V) I, \tag{25}$$
$$C' + O + I = 1,$$

where we also note that we return to the original definition of $O$.

##### 1.2.7 Iteration 7

The seventh iteration corresponded to the boundary where $p_{10} \to 0$. As this parameter appears in the rate $p_9 \exp(-p_{10} V)$, we can simply write this rate as $p_9$, giving the new system equations:

$$\dot{O} = \phi_5 \exp(p_{12} V) C' + p_9 I - (\phi_6 \exp(-p_4 V) - p_7 \exp(p_8 V)) O \tag{26}$$

$$\dot{I} = p_7 \exp(p_8 V) O - p_9 I, \tag{27}$$

with

$$C' + O + I = 1.$$

###### 1.2.8 Iteration 8

The eighth iteration corresponded to the boundary where $p_7, p_9 \to \infty$. Dividing Equation 26 by $p_9$ gives us

$$I = \phi_7 \exp(p_8 V) O,$$

where $\phi_7 = p_7/p_9$. Due to the fact that O can be expressed as a function of I, we can consider a new state, $O''$, which is equal to $O + I$. This gives us

$$\dot{O}'' = \phi_5 \exp(p_{12} V) C' - \phi_6 \exp(-p_4 V) O,$$

where

$$I = \frac{\phi_7 \exp(p_8 V)}{1 + \phi_7 \exp(p_8 V)} O'',$$
$$O = \frac{O''}{1 + \phi_7 \exp(p_8 V)}.$$

We are thus left with a single system equation, given by

$$\dot{O}'' = \phi_5 \exp(p_{12} V)(1 - O'') - \frac{\phi_6 \exp(-p_4 V)}{1 + \phi_7 \exp(p_8 V)} O'', \tag{28}$$

which we solve as a single ODE but note that the actual open probability is given by $O = O''/(1 + \phi_7 \exp(p_8 V))$.

###### 1.2.9 Iteration 9

The ninth iteration corresponded to the boundary where $\phi_6, \phi_7, p_{15} \to \infty$, where $p_{15}$ is the maximal conductance parameter. From this we can write $\phi_6 \exp(-p_4 V)/(1 + \phi_7 \exp(p_8 V))$ as $\psi_6 \exp(-(p_4 + p_8)V)$ where $\psi_6 \approx \phi_6/\phi_7$ and we can write $p_{15}/(1 + \phi_7 \exp(p_8 V))$ as $\phi_{15} \exp(-p_8 V)$ where $\phi_{15} \approx p_{15}/\phi_7$. We are thus left with a single system equation, given by

$$\dot{O}'' = \phi_5 \exp(p_{12} V)(1 - O'') - \psi_6 \exp(-(p_4 + p_8)V) O'', \tag{29}$$

with the following expression for the current:

$$I_{\mathrm{Kr}} = \phi_{15} \exp(-p_8 V) \cdot O'' \cdot (V - E_{\mathrm{Kr}}). \tag{30}$$

### 2 Supplementary Figures

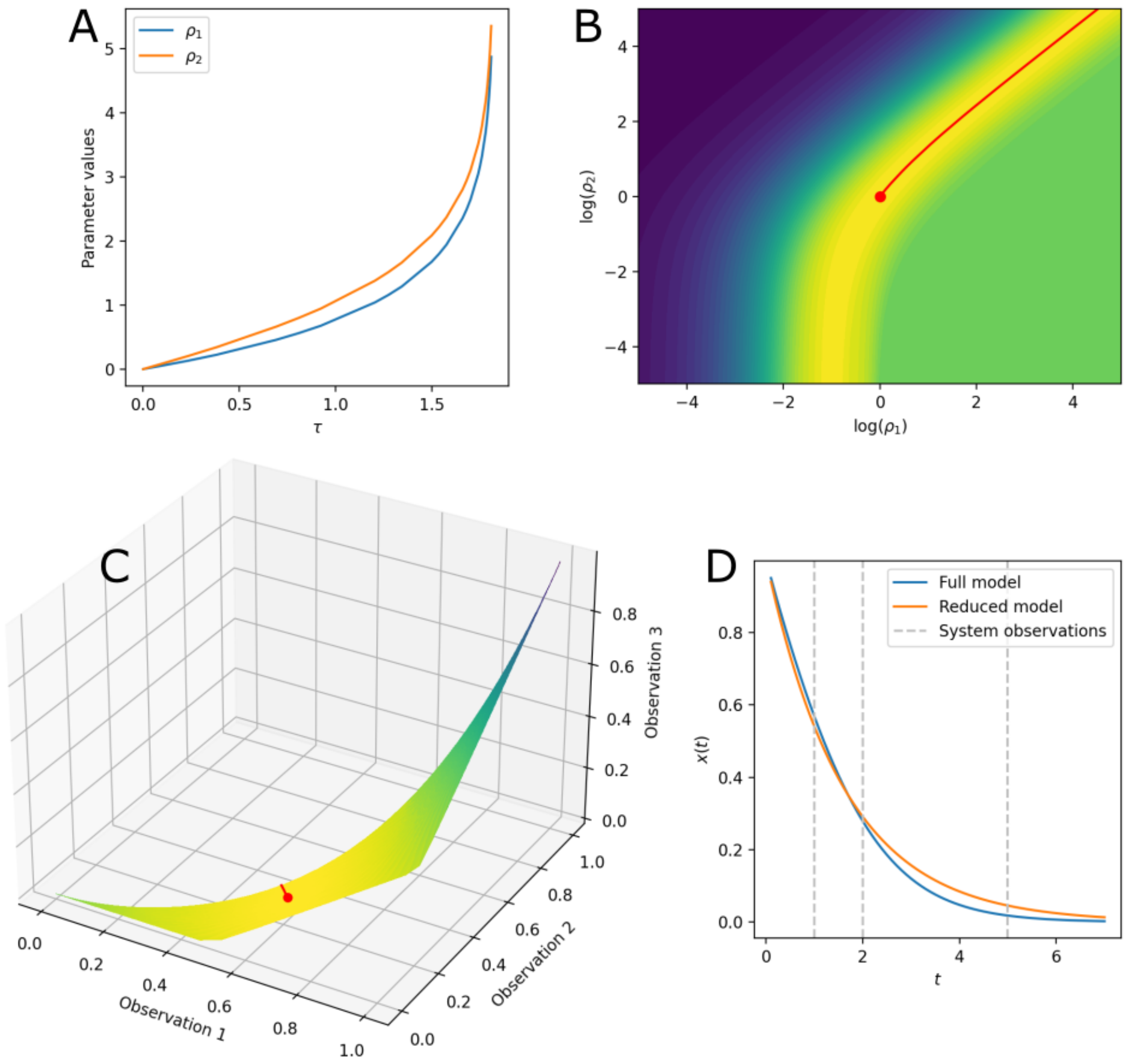


Figure S1: Application of the manifold boundary approximation method to a Michaelis Menton reaction kinetics "toy model", adapted from `https://github.com/mktranstrum/MBAM`. (A) Parameter values along the geodesic path. The geodesic path in (B) parameter space and (C) along the model manifold, from the initial position denoted by a circle, coloured according to the cost function value such that yellow represents a good fit and green/blue represent worse fits. (D) A comparison of the full and reduced models, given by Equations 1 and 2, respectively, and the times at which they are observed.

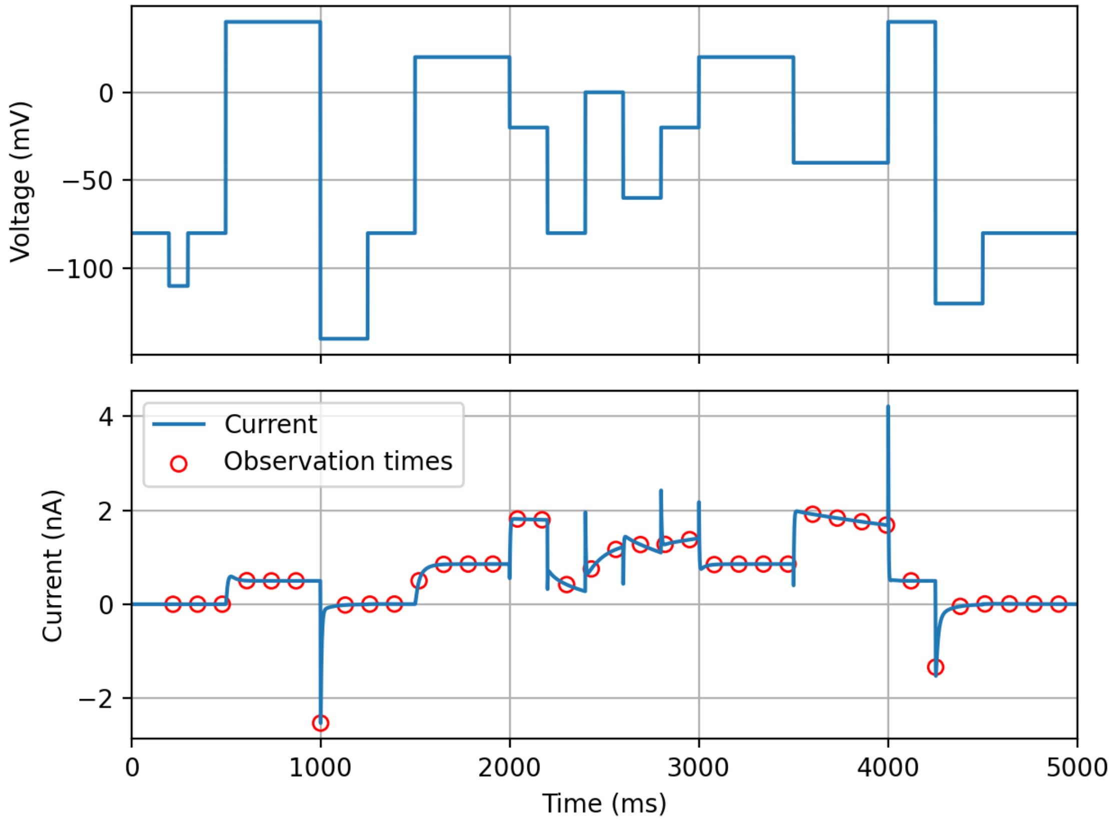


Figure S2: (Upper panel) the input voltage protocol used for iterations of the manifold boundary approximation method. (Lower panel) observed current for the full hERG channel model (blue line) and times at which observations are made (red circles).

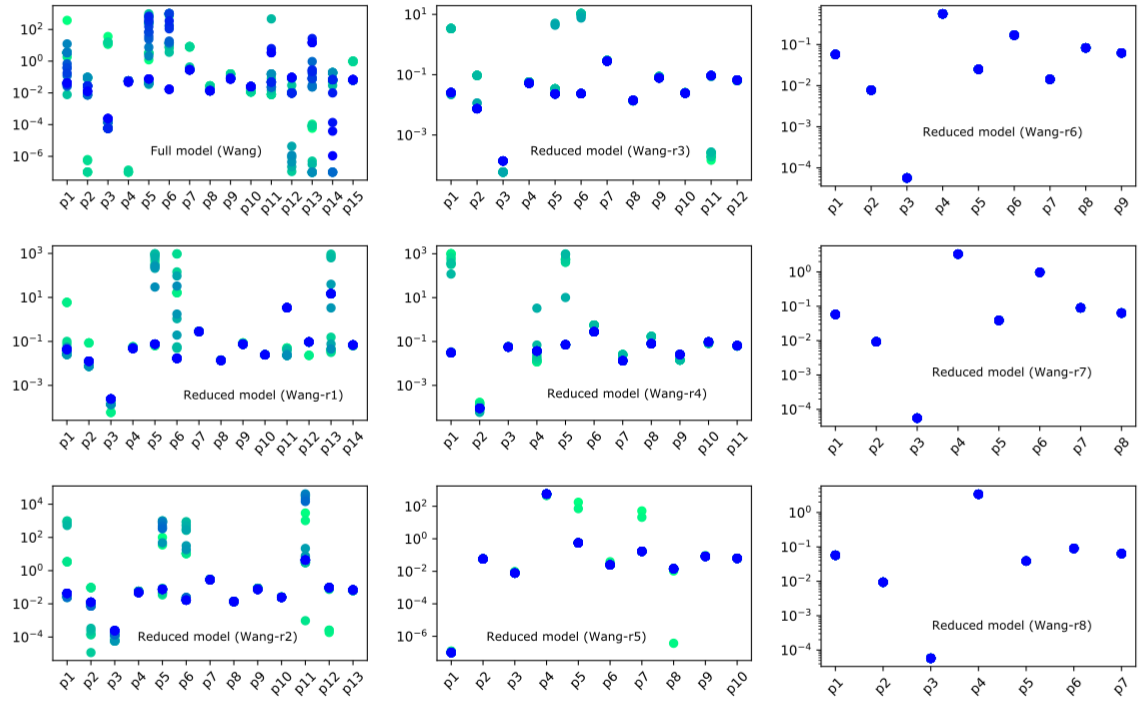


Figure S3: Inferred parameter values for the full Wang model [3] and a series of reduced models for 50 repeats of fitting from different initial guesses to experimental 'staircase' calibration protocol hERG channel currents at 37°C [1]. The 30 best parameter sets are shown in each case, from lowest (green) to highest (blue) likelihood.

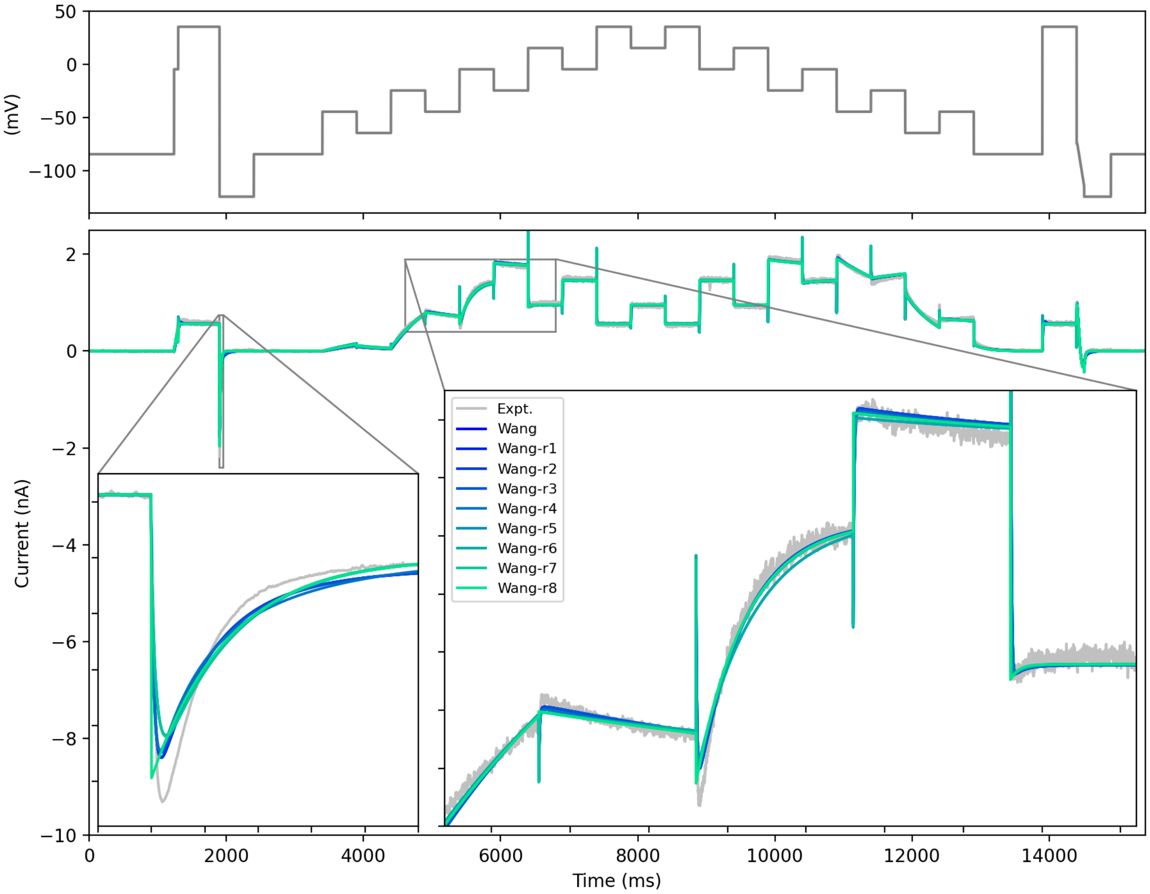


Figure S4: A comparison of the full Wang model and series of reduced model fits to experimental data under the 'staircase' calibration protocol.

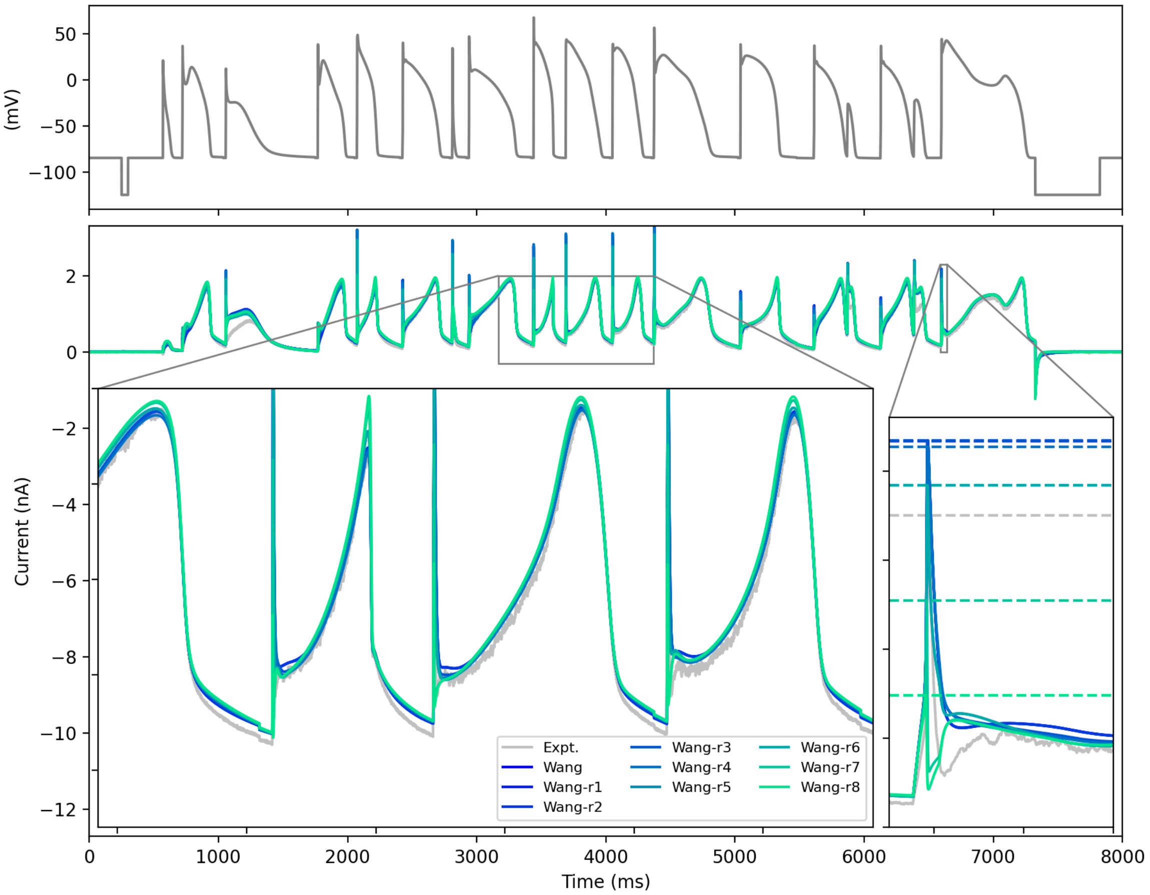


Figure S5: Predictions of the full Wang model and series of reduced models under a complex action potential waveform validation protocol. The dotted lines in the lower right inset show the peaks of the hERG1a transient, coloured according to the model or experiment (grey).

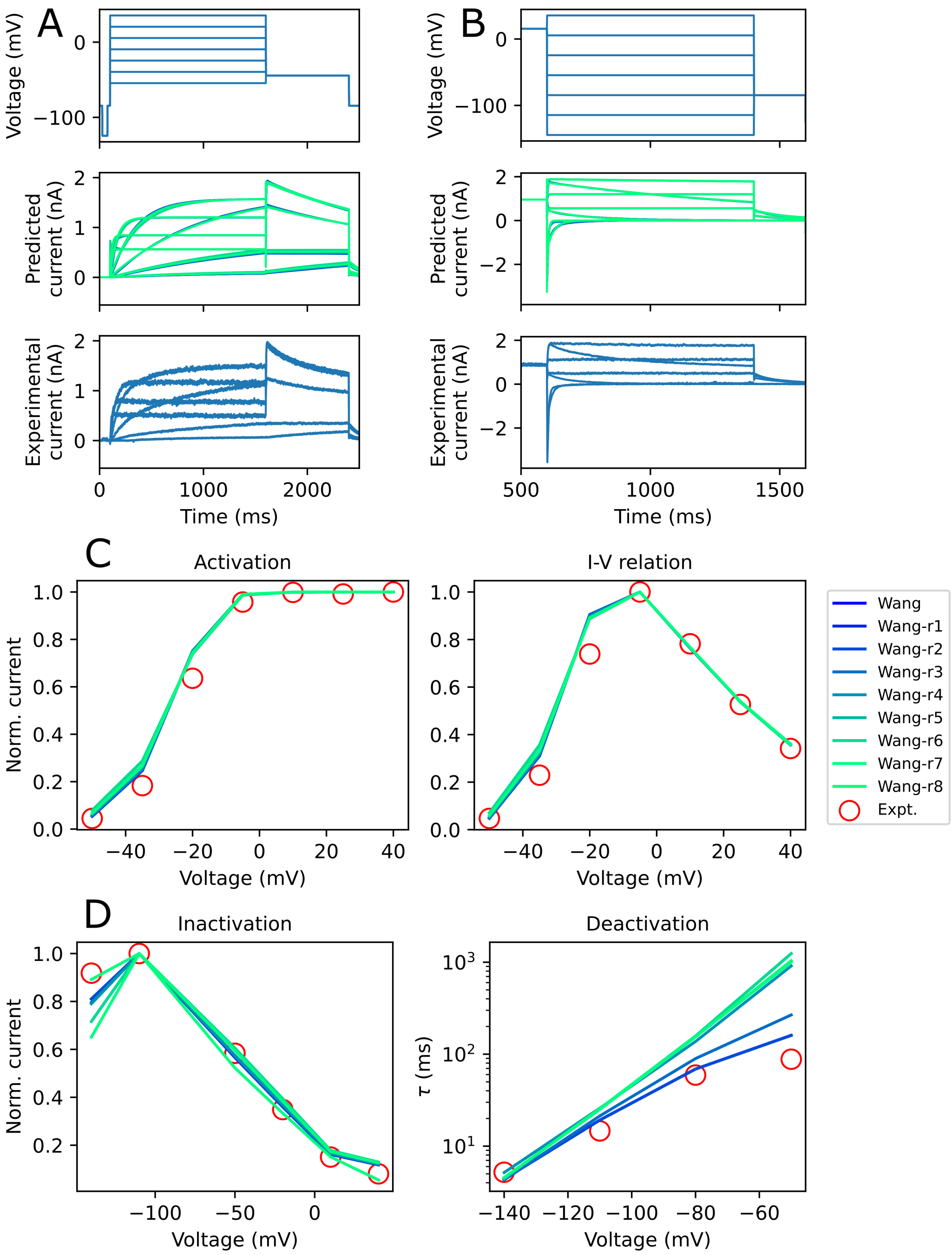


Figure S6: Predictions of the full Wang model and series of reduced models under shortened versions of traditional (A) activation and (B) inactivation protocols. Comparisons of summary data between the full Wang model and reduced models for (C) activation and (D) inactivation data, corresponding to the experiments shown in (A,B). All experimental data were recorded in HEK293 cells at 37°C.
